# Eosinophil-derived COX-2 protects against experimental colitis through the PGE_2_-IL-22 axis

**DOI:** 10.64898/2025.12.18.695257

**Authors:** Yang Yang, Constance L. Atkins, Yuanyuan Fan, Lidan Hou, Jie Zhao, Junda Gao, Yankai Wen, Nicolas F. Moreno, Xiangsheng Huang, Faraz Bishehsari, Agnieszka Czopik, Sean P. Colgan, Tugrul Purnak, Keith C. Summa, Yingzi Cong, Elizabeth A. Jacobsen, Cynthia Ju

## Abstract

Inflammatory bowel disease (IBD) is driven by a breakdown in immune regulation and epithelial barrier function, yet the contribution of eosinophils to this process has remained poorly defined and controversial. While eosinophils infiltrate the intestinal mucosa during both flares and remission, their role in shaping disease outcomes is unclear. Our RNA-seq analyses of colonic eosinophils isolated from dextran sulfate sodium (DSS)-treated mice revealed a significant upregulation of cyclooxygenase (Cox)-2 (gene name, *Ptgs2*). Eosinophil-specific deletion of Cox-2 (Ptgs2^fl/fl^eoCre^+/-^) reduced IL-22 production and exacerbated DSS- and trinitrobenzene sulfonic acid (TNBS)-induced colitis, characterized by greater weight loss, higher disease activity, colon shortening, and epithelial injury. Administration of recombinant IL-22 reversed these phenotypes. Mechanistically, eosinophil-derived COX-2 enhanced IL-22 production by type 3 Innate lymphoid cells (ILC3s) through prostaglandin E2 (PGE_2_) signaling. Consistently, Ptgs2^fl/fl^eoCre^+/-^mice exhibited reduced colonic PGE_2_ levels, while PGE_2_ analog treatment restored IL-22 production and mucosal protection. Our findings identify eosinophil-derived COX-2 and PGE_2_ as a critical regulator of IL-22 production during colitis, uncovering a previously unrecognized eosinophil-ILC3 crosstalk that safeguards the intestinal barrier and represents a promising therapeutic target in IBD.

**Significance Statement:** The role of eosinophils in inflammatory bowel disease (IBD) has remained unresolved. We identify eosinophil-derived COX-2 as the main source of prostaglandin E2 (PGE_2_) in colitis, which drives IL-22 production by ILC3s to maintain epithelial protection. This discovery defines a protective eosinophil/PGE_2_/IL-22 circuit and reveals a therapeutic target to strengthen barrier defense in IBD.

## Introduction

Inflammatory bowel disease (IBD), comprising Crohn’s disease (CD) and ulcerative colitis (UC), is characterized by chronic inflammation, recurrent flares, and disruption of the intestinal epithelial barrier(1, 2). Innate immune cells play central roles in IBD, exerting both protective and pathogenic functions depending on the context(3). For example, neutrophils release reactive oxygen species (ROS), cytotoxic granules and neutrophil extracellular traps (NETs) that exacerbate tissue damage(4-7), yet they are also crucial for resolving inflammation, facilitating epithelial repair, and promoting mucosal healing (8-11).

Eosinophils are reported to accumulate in the inflamed intestinal mucosa of IBD patients(12, 13), but their precise role in the disease pathogenesis and protection remains poorly understood. Clinical studies report conflicting associations: increased eosinophil abundance and granule protein release correlate with more severe disease in some cohorts(12, 14, 15), while other reports link higher eosinophil counts to remission in UC(16, 17). Notably, patients responding to therapies such as Ustekinumab and Vedolizumab often display elevated eosinophil counts(18), and eosinophil-predominant inflammation has been associated with a reduced risk of flares compared to neutrophil-predominant disease(19). There are very few pre-clinical studies in the literature investigating the role of eosinophils in IBD, and they have yielded contradictory results. In IL-10^-/-^ mice, eosinophil depletion by CCR3 deletion did not alter colitis severity(20), whereas antibody-mediated eosinophil depletion reduced dextran sodium sulfate (DSS)- induced colitis(21). By contrast, eosinophil-deficient (PHIL) mice exhibited worsened DSS colitis due to the loss of anti-inflammatory lipid mediators(22). These discrepancies underscore that context-dependent functions of eosinophils in intestinal inflammation and highlight the need to clarify their mechanistic contributions.

IL-22 has emerged as a key cytokine in maintaining epithelial integrity(23). Deficiency in IL-22 exacerbates colitis, while exogenous administration ameliorates disease in mouse models(24, 25). IL-22 signals through intestinal epithelial cells (IECs) to activate STAT3, stimulate proliferation, and enhance production of mucins and antimicrobial peptides, thereby reinforcing the epithelial barrier(26-29). Type 3 Innate lymphoid cells (ILC3s) and T helper cells (Th17, Th22) represent the primary sources of IL-22, typically induced by cytokines such as IL-23 and IL-1β(30-32). Additional stimuli, including TLR2 agonists, aryl hydrocarbon receptor (AHR) ligands, and NK cell receptor engagement also regulate IL-22 production(33-35). However, the role of other immune cell types in orchestrating IL-22 responses during colitis remains incompletely understood.

Here, we identify eosinophils as a critical regulator of IL-22 production in experimental colitis. We show that eosinophils are a major source of cyclooxygenase-2 (COX-2) in the inflamed colon and that eosinophil-derived COX-2 drives PGE_2_ synthesis, which enhances IL-22 secretion by ILC3s. Using eosinophil-specific Cox-2-deficient mice, we uncovered a previously unrecognized eosinophil-PGE_2_-IL-22 axis that mitigates intestinal inflammation. These findings clarify the protective role of eosinophil-ILC3 circuit in colitis and highlight the COX2-PGE_2_-IL-22 pathway as a promising therapeutic target for colitis.

## Result

### Eosinophil-derived COX-2 plays a protective role during experimental colitis

Although eosinophils are recognized for their roles in immune regulation and tissue remodeling(36), their function in colitis remains unclear. To address this, we analyzed gene expression profiles of eosinophils in a mouse model of DSS-induced colitis (DSS-colitis). Colonic eosinophils were isolated from water (control)- and DSS-treated C57Bl/6 WT mice and subjected to bulk RNA sequencing. Pathway analyses using KEGG and DAVID identified 40 differentially expressed genes, which were validated by qPCR (**Fig.1A-B, Fig.S1)**. Among these, Ptgs2 (the gene encoding cyclooxygenase-2, Cox-2) was the most significantly upregulated gene in colonic eosinophils from DSS-treated mice compared with controls (**Fig.1B**). Consistent with this finding, Ptgs2 mRNA levels in whole colonic tissue were markedly elevated following DSS exposure (**Fig.1C**).

**Fig. 1.**
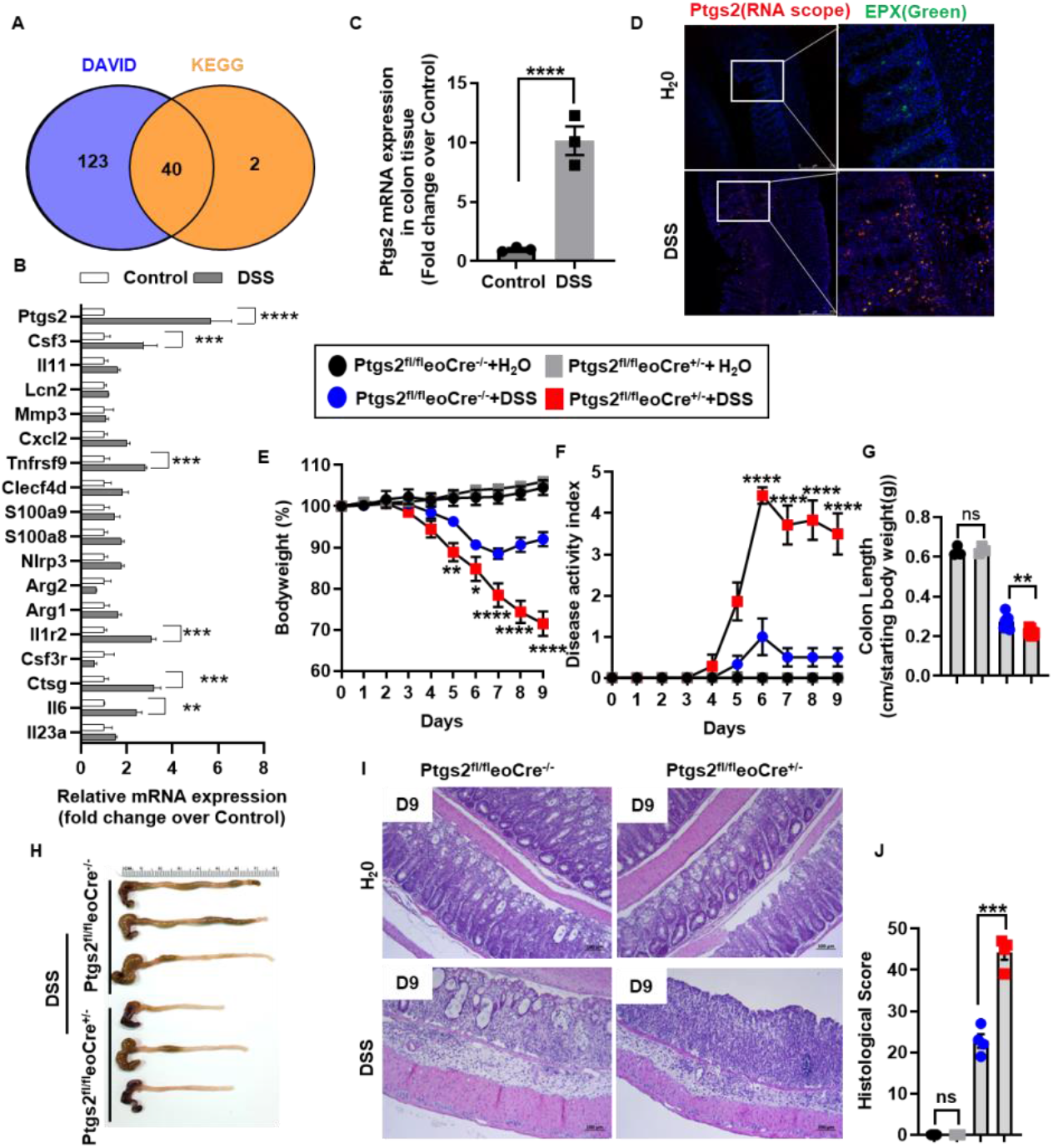
Deletion of Ptgs2 in eosinophils renders more severe colitis. (A-D) WT C57Bl/6 mice were treated with regular water or 3% DSS for 5 days. On day 6, colonic eosinophils were isolated/purified by MACS and subjected to RNAseq and qPCR analyses. (n=3/group). (A) 40 DEGs were identified by KEGG/DAVID pathway analyses of RNAseq data. (B) mRNA levels of DEGs in colonic eosinophils were measured q-PCR. (C) Ptgs2 mRNA levels in colon tissue were measured by q-PCR. (D) RNAscope analysis of Ptgs2 and immunofluorescence staining for EPX expression in the mouse colon. (E-J) Ptgs2^fl/fl^eoCre^+/-^ mice and their WT littermates (Ptgs2^fl/fl^eoCre^-/-^) were treated with 3% DSS for 5 day and allowed to recover for 4 days before collection of colon tissues on day 9. (n=4/group) (E) Body weight changes (F) disease activity index and (G-H) colon lengths were measured. (I-J) Representative colon histology and histological scores. Data are means ± SEM. In all scatter plots with bars, each data point represents one individual. A two-tailed unpaired Student’s t test with Welch’s correction was performed in B and C. Two-way ANOVA was performed in E, F, G, and J. *p < 0.05, **p < 0.01, ***p < 0.001, ****p < 0.0001. All experiments were repeated a minimum of three times.

Because Cox-2 can also be expressed by epithelial cells(37, 38), macrophages(39, 40), and endothelial cells(41, 42), we next determined which cell type was the predominant source during colitis. Using RNAscope to detect Ptgs2 mRNA in combination with immunofluorescence (IF) staining for eosinophil peroxidase (EPX), we observed a significant increase in Ptgs2-expressing cells in DSS-treated colon tissues, with the majority of these cells identified as EPX^+^ eosinophils (**Fig.1D**). These data indicate that eosinophils are the major source of Ptgs2 in the inflamed colon.

To directly investigate the functional role of eosinophil-derived COX-2 in colitis, we generated eosinophil-specific Cox-2-deficient mice (Ptgs2^fl/fl^eoCre^+/-^). Compared with DSS-treated WT littermates (Ptgs2^fl/fl^eoCre^-/-^), Ptgs2^fl/fl^eoCre^+/-^ mice exhibited significantly greater weight loss (**Fig.1E**), elevated disease activity index (DAI) scores (**Fig.1F**), shortened colons (**Fig.1G-H**), and more severe histopathology, including increased inflammatory infiltration, epithelial damage, and crypt loss (**Fig.1I-J**). To confirm these findings, we employed a second model of experimental colitis induced by 2,4,6-trinitrobenzene sulfonic acid (TNBS). Similar to the DSS model, TNBS-treated Ptgs2^fl/fl^eoCre^+/-^ mice developed more severe disease than WT littermates, as evidenced by greater weight loss (**Fig.S2A**), shorter colon length (**Fig.S2B**) and exacerbated histological inflammation (**Fig.S2C-D**). Together, these results demonstrate that eosinophil-derived COX-2 plays a protective role in acute colitis.

### Eosinophil-specific deletion of Cox-2 reduces colonic IL-22 production

To understand how eosinophil-derived COX-2 protects against DSS-colitis, we first examined whether Cox-2 deletion alters eosinophil abundance in the colon. No significant differences were observed under homeostasis or after DSS treatment (**Fig.2A-B**), indicating that COX-2 does not affect eosinophil recruitment or survival.

Because eosinophils are known to modulate immune responses through cytokine regulation(43, 44), for example, by suppressing Th17 cells via constitutive IL-1Ra secretion(45), we next asked whether eosinophil-derived COX-2 influences cytokine production during colitis. ELISA of colon tissues on day 6 after DSS treatment revealed higher levels of IL-1β and IL-6 in Ptgs2^fl/fl^eoCre^+/-^ mice compared with WT littermates, consistent with more severe inflammation (**Fig.S3**). Strikingly, both mRNA and protein levels of IL-22 were dramatically reduced in Ptgs2^fl/fl^eoCre^+/-^ colons (**Fig.2C and Fig. S3**). *Ex vivo* cultures confirmed a marked reduction in IL-22 release from Ptgs2^fl/fl^eoCre^+/-^ colons (**Fig.2D**).

IL-22 promotes epithelial protection by engaging its receptor and activating STAT3 signaling in intestinal epithelial cells(28, 46). In line with reduced IL-22, STAT3 activation (p-STAT3) was significantly diminished in DSS-treated Ptgs2^fl/fl^eoCre^+/-^ mice compared with WT controls (**Fig. 2E-F**). To test whether IL-22 deficiency accounted for worsened disease in Ptgs2^fl/fl^eoCre^+/-^ mice, we administered recombinant IL-22 (rmIL-22) during DSS treatment. Remarkably, exogenous rmIL-22 alleviated colitis severity, reducing epithelial injury, accelerating weight recovery, improving DAI scores, and restoring colon length (**Fig. 2G-L**). These findings demonstrate that eosinophil-derived COX-2 protects against DSS-colitis by sustaining IL-22 production and downstream epithelial protective signaling.

**Fig. 2.**
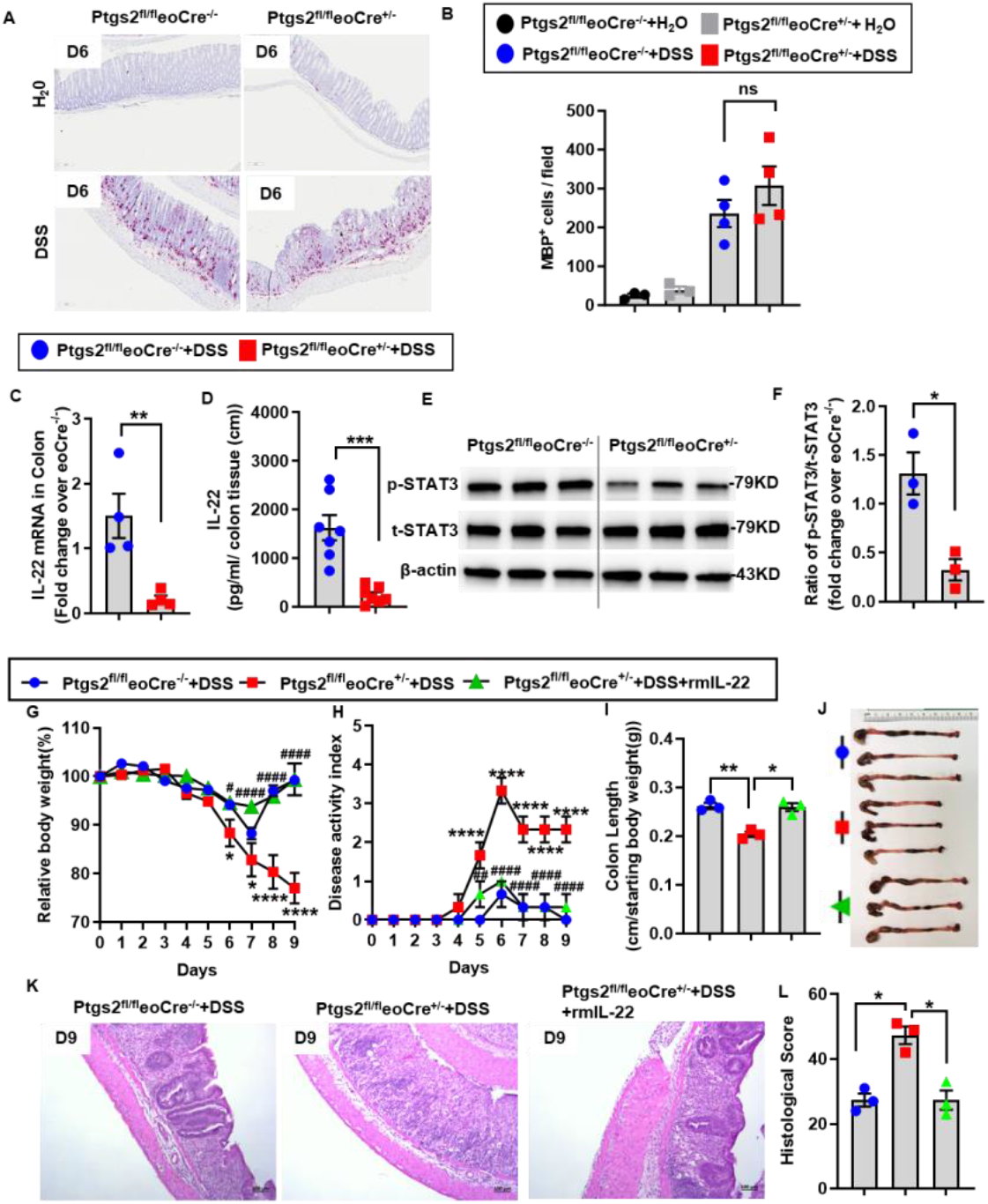
Deletion of Ptgs2 in eosinophils impaired IL-22 production. (A-F) Ptgs2^fl/fl^eoCre^+/-^ mice and their WT littermates were treated with 3% DSS for 5 days. Colon tissues and IECs were collected on day 6. The control mice group was treated with regular water. (A-B) IHC staining for eosinophils by anti-mouse major basic protein (MBP) antibody, and the numbers of MBP^+^ cells were quantified. (n=4/group) (C) mRNA levels of IL-22 in colon tissue were measured by q-PCR (n=4/group). (D) Protein levels of IL-22 were measured in the colon culture supernatant. (E-F) p-STAT3 and t-STAT3 protein expression were detected by Western blotting and quantified. (n=3/group) (G-L) Ptgs2^fl/fl^eoCre^+/-^ mice and their WT littermates were treated with 3% DSS for 5 days and allowed to recover for 4 days before collection of colon tissues on day 9. rmIL-22 was given on days 2, 5, and 8 of DSS treatment. (n=3/group) (G) Body weight changes, (H) disease activity index and (I-J) colon lengths were measured. (K-L) Representative colon histology and histological scores. Data are means ± SEM. In all scatter plots with bars, each data point represents one individual. A two-tailed unpaired Student’s t test with Welch’s correction was performed in C, D, and F. Two-way ANOVA was performed in B, G, H, I, and L. *p < 0.05, **p < 0.01, ***p < 0.001, ****p < 0.0001; ^#^p < 0.05, ^##^p < 0.01, ^###^p < 0.001, ^####^p < 0.0001. All experiments were repeated a minimum of three times.

### Eosinophil-derived PGE_2_ plays a critical role in IL-22 production by colonic ILC3 cells

COX-2 is a key enzyme in the biosynthesis of prostaglandins (PGs), including PGE_2_, PGD_2_, PGF_1α_, and PGF_2α_ (47). To determine whether eosinophil-derived COX-2 contributes to prostaglandin production during colitis, we measured PGs in colonic tissues of DSS-treated Ptgs2^fl/fl^eoCre^+/-^ and WT littermates using LC/MS. PGE_2_ levels were significantly lower in Ptgs2^fl/fl^eoCre^+/-^ than in WT controls (**Fig.3A**), a finding confirmed by ELISA (**Fig.3B and Fig.S4**).

To test whether loss of PGE_2_ accounted for reduced IL-22 and worsened DSS-colitis, we treated Ptgs2^fl/fl^eoCre^+/-^ mice with a stable PGE_2_ analog, 16, 16-dimethyl prostaglandin E2 (dmPGE_2_). Remarkably, dmPGE_2_ treatment ameliorated DSS colitis in mutant mice, restoring disease severity to levels comparable to WT littermates (**Fig. 3C-H**). Importantly, dmPGE_2_ also rescued colonic IL-22 production (**Fig.3I**). We next examined the direct impact of PGE_2_ on IL-22 production. *Ex vivo* cultures of colon tissues stimulated by IL-23 showed that addition of dmPGE_2_ significantly enhanced IL-22 secretion. This effect was mediated specifically via the EP4 receptor, as EP4 inhibition, but not blockade of other EP receptors, abrogated the response (**Fig. 3J**).

**Fig. 3.**
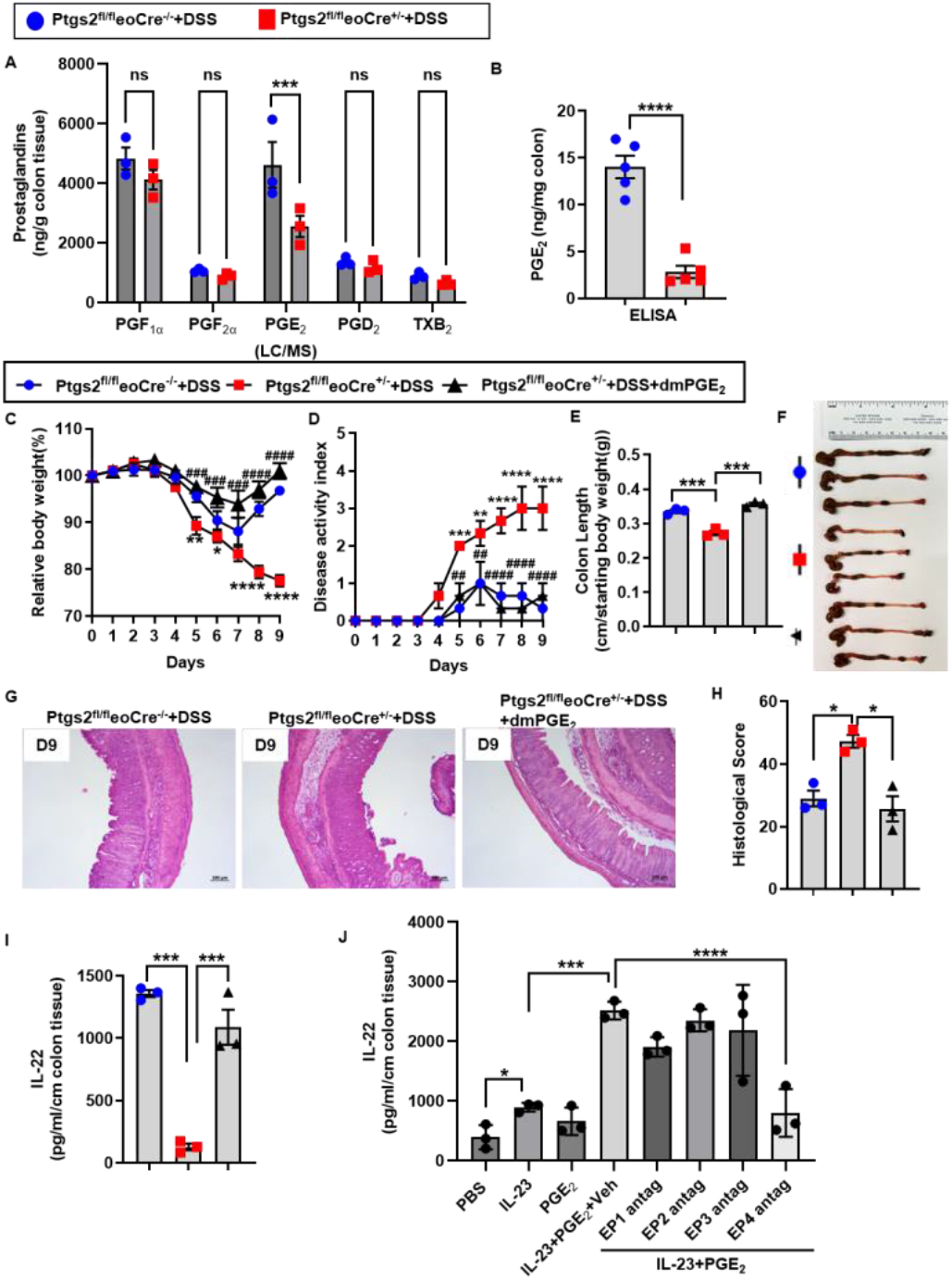
Eosinophil mediates IL-22 production through PGE_2_/EP4 signaling. (A-B) Ptgs2^fl/fl^eoCre^+/-^ mice and their WT littermates were treated with 3% DSS for 5 days. Colon tissues were collected on day 6. (A) Colonic PGF_1α_, PGF_2α_, PGE_2_, PGD_2_, and TXB_2_ levels were measured by LC/MS(n=3/group). (B) Colonic PGE_2_ levels were measured by ELISA (n=5/group). (C-I) Ptgs2^fl/fl^eoCre^+/-^ mice and their WT littermates were treated with 3% DSS for 5 day and allowed to recover for 4 days before collection of colon tissues on day 9. dmPGE_2_ was given on days 2, 5, and 8 of DSS treatment. (n=3/group) (C) Body weight changes, (D) disease activity index and (E-F) colon lengths were measured. (G-H) Representative colon histology and histological scores. (I) Protein levels of IL-22 were measured in the supernatant. (J) Colon tissues were collected on D6 after DSS treatment of WT mice and then cultured with IL-23, dmPGE_2,_ and different PGE_2_ receptor antagonists for 2 days. Protein levels of IL-22 were measured in the colon culture supernatant. Data are means ± SEM. In all scatter plots with bars, each data point represents one individual. A two-tailed unpaired Student’s t test with Welch’s correction was performed in B. Two-way ANOVA was performed in A, C, D, E, H, I and J. *p < 0.05, **p < 0.01, ***p < 0.001, ****p < 0.0001; ^#^p < 0.05, ^##^p < 0.01, ^###^p < 0.001, ^####^p < 0.0001. All experiments were repeated a minimum of three times.

IL-22 is produced by multiple intestinal immune subsets, including ILC3s, Th17 cells, γδ T cells, and Treg cells(32, 48-51). Our flow cytometric analyses revealed that CCR6-expressing Lymphoid Tissue Inducer-like cells (CCR6^+^ LTi-like ILC3s; CD45^+^Lineage^-^ CD90.2^+^RORγt^+^CCR6^+^) represented the predominant source of IL-22 in colons of DSS-treated WT mice (**Fig. S5A**). Recent single-cell RNA-sequencing analyses of the healthy mouse gut(52), we also measured another two principal subsets within CD90.2^+^RORγt^+^ ILC3s in colonic lamina propria by flow cytometry: T-bet^+^ (NKp46^+^) ILC3s and CCR6^−^T-bet^−^ “double-negative” (DN) ILC3s. Compared to CCR6^+^ LTi-like ILC3s, NKp46^+^ ILC3s produce comparatively lower levels of IL-22 and DN ILC3s show undetectable IL-22 expression (**Fig.S5B-D**). While Ptgs2 deletion in eosinophils did not alter ILC3 total number and their subsets abundance (**Fig. 4A-C and Fig.S5E-G**), it markedly reduced the proportion of IL-22^+^CCR6^+^ILC3s (**Fig. 4D-F and Fig.S5B-D**). Moreover, treating isolated lamina propria lymphocytes with dmPGE_2_ significantly increased IL-22^+^ ILC3s, an effect abolished by EP4 antagonism (**Fig. 4D–F**). Together, these findings identify eosinophil-derived COX-2 as a critical upstream regulator of IL-22 production by colonic ILC3s through the COX-2-PGE_2_-EP4 axis.

**Fig. 4.**
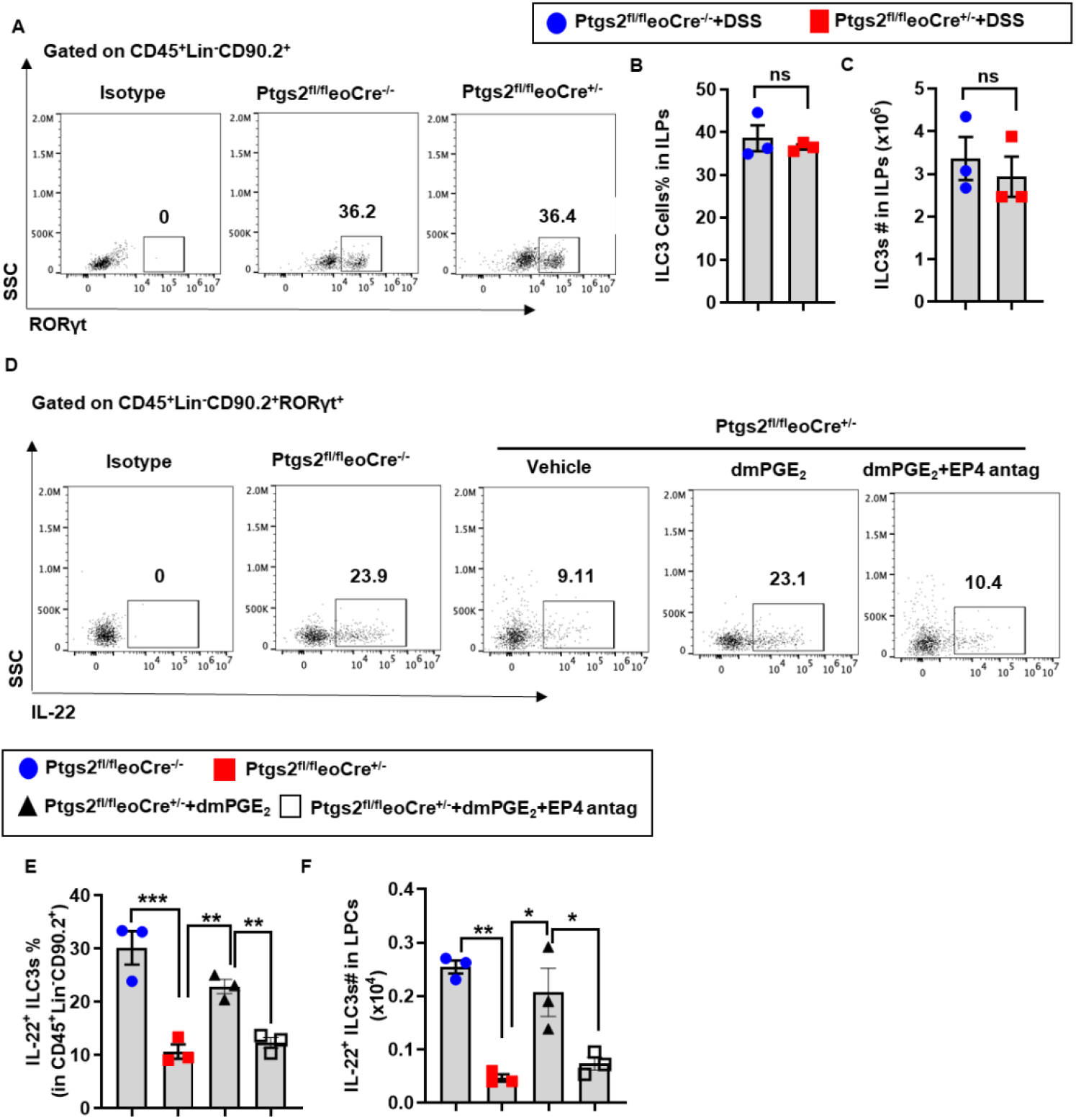
Eosinophil-derived PGE2 is critical for CCR6^+^ILC3s to produce IL-22. (A-C) Ptgs2^fl/fl^eoCre^+/-^ mice and their WT littermates were treated with 3% DSS for 5 days. On day 6, colon lamina propria cells (LPs) were isolated and divided into two groups. One group was directly stained for intracellular RORγt to detect ILC3 numbers; the other group was cultured with dmPGE_2_ or EP4 antagonists for intracellular IL-22 staining (n=3/group) (A-C) Representative flow cytometry, the percentage and numbers of ILC3s (CD45^+^Lin^-^ CD90.2^+^RORγt^+^) in the LPs were measured and quantified. (D-F) Representative flow cytometry, the percentage and numbers of IL-22^+^ILC3s (CD45^+^Lin^-^CD90.2^+^RORγt^+^IL-22^+^) in the LPs were measured and quantified. Data are means ± SEM. In all scatter plots with bars, each data point represents one individual. A two-tailed unpaired Student’s t test with Welch’s correction was performed in B and C. Two-way ANOVA was performed in E and F. *p < 0.05, **p < 0.01, ***p < 0.001. All experiments were repeated a minimum of three times.

### The PGE_2_-IL-22 axis mediates eosinophil-driven protection in colitis

A previous study using eosinophil-deficient (PHIL) mice demonstrated that eosinophils protect against DSS-colitis by producing the anti-inflammatory lipid protectin D1(22). To validate the protective function of eosinophils, we employed a separate eosinophil-deficient strain (ΔdblGATA1) and confirmed that eosinophil deficiency exacerbated DSS-colitis (**Fig. S6**). Importantly, ELISA revealed a significant reduction in colonic IL-22 levels in ΔdblGATA1 mice compared with WT controls (**Fig. 5F**), a phenotype closely resembling that observed in Ptgs2^fl/fl^eoCre^+/-^ mice (**Fig.2G-I**).

**Fig. 5.**
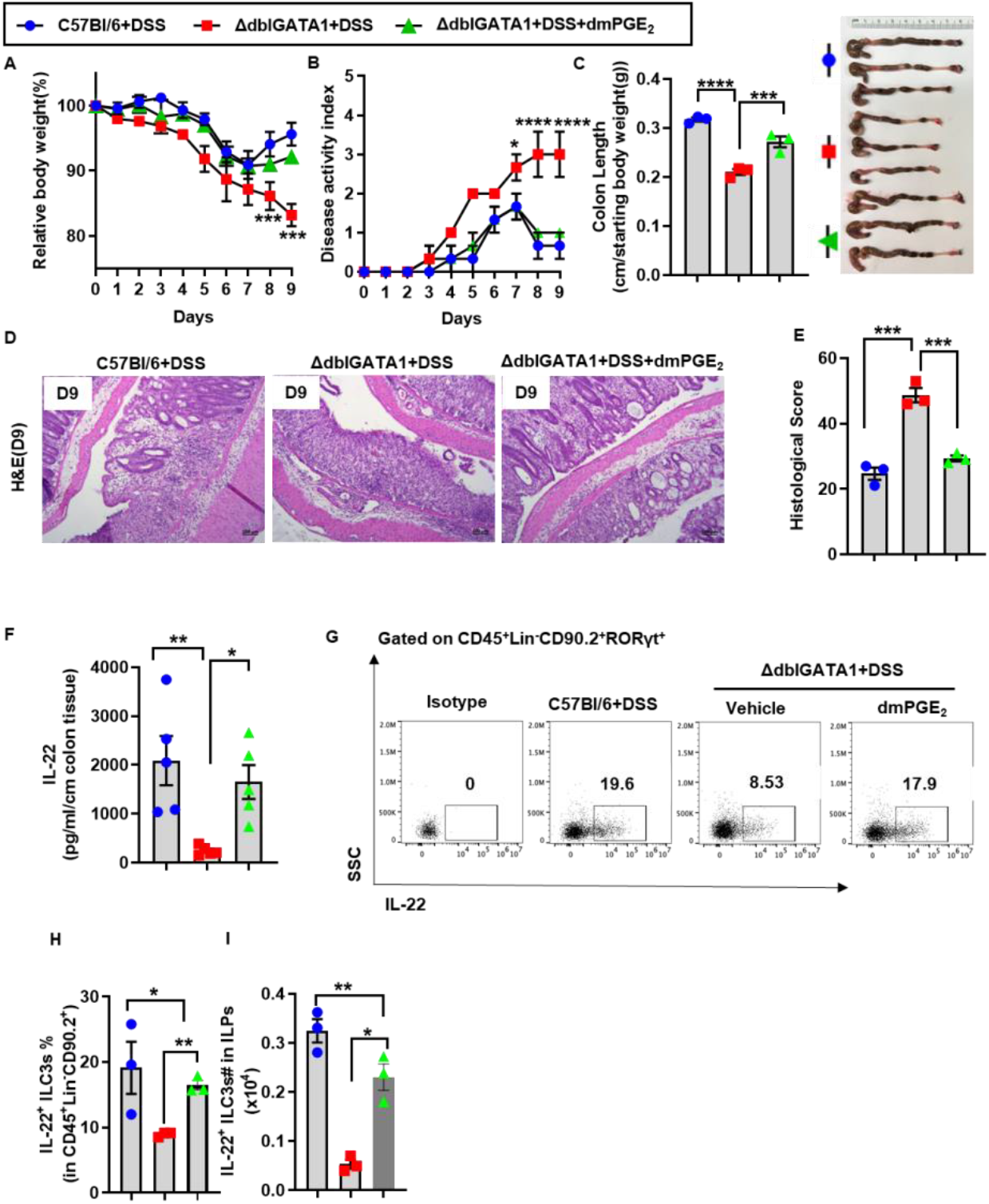
The PGE2-IL-22 axis mediates eosinophil-driven protection in colitis. (A-E) ΔdblGATA1 mice and their WT counterparts were treated with 3% DSS for 5 days and allowed to recover for 4 days before the collection of colon tissues on day 9. (F-I) Colon tissue and lamina propria cells (LPs) were isolated on D6. dmPGE_2_ was given on days 2, 5, and 8 of DSS treatment. (n=3/group) (A) Body weight changes, (B) disease activity index and (C) colon lengths were measured. (D-E) Representative colon histology and histological scores. (F) Protein levels of IL-22 were measured in the supernatant. (G-I) Representative flow cytometry, the percentage and numbers of IL-22^+^ILC3s (CD45^+^Lin^-^ CD90.2^+^RORγt^+^IL-22^+^) in the LPs were measured and quantified. Data are means ± SEM. In all scatter plots with bars, each data point represents one individual. Two-way ANOVA was performed in A, B, C, E, F, H, and I. *p < 0.05, **p < 0.01, ***p < 0.001, ****p < 0.001. All experiments were repeated a minimum of three times.

To determine if the PGE_2_-IL-22 axis underlies this phenotype, we treated ΔdblGATA1 mice with dmPGE_2_ during DSS administration. dmPGE_2_ treatment markedly alleviated disease severity, as shown by accelerated weight recovery, reduced DAI scores, restoration of colon length, and decreased epithelial damage (**Fig. 5A-E**). Notably, dmPGE_2_ also restored IL-22 protein levels toward those of WT mice (**Fig. 5F**). Finally, to determine whether ILC3s in ΔdblGATA1 mice retain the capacity to produce IL-22, we analyzed colonic lamina propria cells following PGE_2_ treatment. Flow cytometric analysis revealed a significant increase in IL-22^+^ ILC3s in ΔdblGATA1 mice after PGE_2_ supplementation (**Fig.5G-I**), indicating that the defect lies in upstream regulation rather than intrinsic dysfunction of ILC3s.

Together, these findings demonstrate that eosinophils promote colonic protection through the PGE_2_-IL-22 axis, positioning them as key upstream regulators of epithelial defense in colitis.

## Discussion

Our study demonstrates that eosinophils protect against DSS- and TNBS-induced colitis through a COX-2-dependent mechanism. Specifically, eosinophil-derived COX-2 drives PGE_2_ production, which in turn promotes IL-22 secretion by ILC3s. This reveals a previously unrecognized eosinophil-ILC3 crosstalk mechanism that protects the epithelium during acute colonic injury.

We identified eosinophils as the predominant source of COX-2 in the inflamed colon. Previous reports suggested macrophages as major COX-2-expressing cells, based on colocalization with markers such as F4/80(41) and CD68(53), whereas other studies failed to confirm these findings(54). These discrepancies highlight the limitations of antibody-based detection. By employing RNAscope to localize Cox2 mRNA, we demonstrated that eosinophils are the major COX-2-expressing population during colitis. This result is consistent with recent single-cell RNA sequencing data showing robust Cox-2-expression in activated intestinal eosinophils(55).

Eosinophil-specific deletion of Ptgs2 markedly exacerbated colonic inflammation, providing the first direct evidence that eosinophil-derived COX-2 is protective in colitis. Interestingly, endothelial cell-derived COX-2 also contributes to protection, primarily via prostacyclin (PGI_2_) (41, 56). However, global deletion of the PGI_2_ receptor does not alter colitis severity(57), suggesting that other prostaglandins may mediate the protective effects. In contrast, epithelial cell-specific Cox-2 deletion does not influence colitis severity(41), despite increased epithelial COX-2 expression after DSS treatment(53). These findings emphasize that the function of COX-2 in colitis is cell-type–dependent, with distinct outcomes determined by both the source of prostaglandins and the responding target cells. This complexity may underlie the conflicting clinical outcomes of COX-2 inhibitor use in IBD patients(58-62) and the concern that such inhibitors may exacerbate gastrointestinal symptoms or trigger flares(63, 64).

Consistent with their role as a source of PGE_2_, eosinophil-specific Ptgs2 deletion significantly reduced colonic PGE_2_ levels and worsened disease severity. Importantly, we show that eosinophils act as upstream regulators of IL-22 production via PGE_2_/EP4 signaling in ILC3s. This mechanism is supported by prior evidence that ILC3s highly express EP4 and that PGE_2_/EP4 signaling enhances IL-23-driven IL-22 production(65). Beyond IL-22, eosinophil-derived PGE_2_ may also stimulate other protective mediators. For example, PGE_2_ can stimulate ILC3s to produce heparin-binding EGF-like growth factor (HB-EGF), which protects epithelial cells from TNF-induced cell death(66). PGE_2_ has been shown to activate macrophages through EP4, enhancing epithelial differentiation and proliferation in DSS-colitis(67). Together, these findings highlight the pleiotropic, cell-type-specific effects of eosinophil-derived PGE_2_ in promoting epithelial protection.

A limitation of this study is the reliance on DSS- and TNBS-induced colitis models, which may not fully capture the complexity of human IBD. While we establish eosinophils as regulators of ILC3-produced IL-22, eosinophil-derived PGE_2_ may also exert direct cytoprotective effects on epithelial cells or modulate other immune populations. The upstream signals that drive Cox-2 induction in eosinophils also remain undefined, although microbial products, TLR4/Myd88 signaling and cytokines such as TNF-α and IL-33 are potential candidates (53, 54, 68, 69).

In conclusion, our work identifies eosinophil-derived COX-2 as a critical regulator of epithelial protection in colitis through the PGE_2_-IL-22 axis. By uncovering eosinophils as a major source of COX-2 and PGE_2_ in the inflamed colon, we define a novel eosinophil-ILC3 interaction that strengthens barrier defense. These insights not only clarify longstanding ambiguities in eosinophil involvement but also point to the PGE_2_-IL-22 pathway as a promising therapeutic target for IBD.

## Materials and Methods

### Animals and experimental colitis mouse model

C57Bl/6J (stock#000664), ΔdblGATA1(on C57Bl/6J background, stock#005653), Ptgs2^flox^ (stock#030785) mice were purchased from the Jackson Laboratory. Breeders of eoCre mice were obtained from Dr. Elizabeth Jacobsen (Mayo Clinic Arizona). Colonies of Ptgs2^fl/fl^eoCre^+/-^, were generated and maintained in the University of Texas Health Science Center at Houston (UTHealth) animal facility. All experiments were performed according to the guidelines of the institutional animal care and use committee (IACUC) at UTHealth. DSS or TNBS-induced colitis models were performed in 10- to 12-week-old male mice as described in *SI Appendix, Supplementary methods*.

### Histology and Immunohistochemistry

Colon tissue sections were fixed in 10% formalin overnight, embedded in paraffin, and cut into 5μm sections. Tissue sections were stained with hematoxylin and eosin (H&E) for the examination of histologic colitis severity. A previously described semi-quantitative scoring system was used to assess histologic colitis severity(70), as described in *SI Appendix, Supplementary methods*.

### RNAscope analysis

RNAscope analysis was performed on paraffin tissue sections using Advanced Cell Diagnostics RNAscope Multiplex Fluorescent Reagent Kit V2 (UM323100), as described in *SI Appendix, Supplementary methods*.

### Isolation of colonic lamina propria cells (LPCs)

LPCs were isolated from the colon using a modified version of a previously described protocol(71), as described in *SI Appendix, Supplementary methods*.

### Flow cytometry analysis

For cell-surface staining, cells (1×10^6^ per tube) were blocked with anti-CD16/CD32 (BioLegend, #101302) and then incubated with the indicated antibodies for 30 min at 4 °C, as described in the *SI Appendix, Supplementary Methods*. Intracellular staining antibodies and procedures are detailed in *SI Appendix, Supplementary Methods*.

### RNA isolation and quantitative real-time polymerase chain reaction (qRT-PCR)

Total RNA was purified from colon tissues or culture cells using Pure-Link RNA Kit (GeneDopt) according to the manufacturer’s protocol, as described in *SI Appendix, Supplementary methods*. 18S rRNA was used as an internal standard. The primers used for qRT-PCR are listed in *SI Appendix, Table S1*.

### *Ex vivo* colon cultures and Enzyme-linked immunosorbent assay (ELISA)

Colon tissues were cultured ex vivo as previously described with minor modifications(72), as described in *SI Appendix, Supplementary methods*

### Statistical analysis

All statistical analyses and graphing were conducted with GraphPad Prism version 10.1.2 (GraphPad Software Inc.). Results were presented as mean ± SEM. For all comparisons in which there were two groups of values, a two-tailed unpaired Student’s t-test with Welch’s correction was performed after demonstrating that the data follow a normal distribution by the Shapiro-Wilk normality test. One-way ANOVA was used to compare values obtained from three or more groups with one independent variable, followed by Tukey’s test. To compare groups with two independent variables, two-way ANOVA was used followed by Tukey’s test. Differences in values were considered significant at *p<0*.*05*. All experiments were repeated a minimum of three times.

## Supporting information

Supplementary Methods, Figures S1 to S6, Tables S1 and SI References

## Acknowledgments

We thank Dr. Feng Li and Dr. Xuan Qin from the NMR and Drug Metabolism Core at Baylor College of Medicine for conducting the LC-MS analysis of prostaglandins. We acknowledge the Mouse Metabolism and Phenotyping Core (MMPC) at the Baylor College of Medicine (BCM, funded by NIH grants R01DK114356 and UM1HG006348).

## Data and materials availability

All data associated with this study are in the paper or the Supplementary Materials.

## References

1. J. Torres, S. Mehandru, J. F. Colombel, L. Peyrin-Biroulet, Crohn’s disease. Lancet 389, 1741–1755 (2017).

2. R. Ungaro, S. Mehandru, P. B. Allen, L. Peyrin-Biroulet, J. F. Colombel, Ulcerative colitis. Lancet 389, 1756–1770 (2017).

3. A. Saez, B. Herrero-Fernandez, R. Gomez-Bris, H. Sanchez-Martinez, J. M. Gonzalez-Granado, Pathophysiology of Inflammatory Bowel Disease: Innate Immune System. Int J Mol Sci 24 (2023).

4. S. Wang et al., Neutrophil-derived PAD4 induces citrullination of CKMT1 exacerbates mucosal inflammation in inflammatory bowel disease. Cell Mol Immunol 21, 620–633 (2024).

5. B. Drury, G. Hardisty, R. D. Gray, G. T. Ho, Neutrophil Extracellular Traps in Inflammatory Bowel Disease: Pathogenic Mechanisms and Clinical Translation. Cell Mol Gastroenterol Hepatol 12, 321–333 (2021).

6. L. A. Denson et al., Clinical and Genomic Correlates of Neutrophil Reactive Oxygen Species Production in Pediatric Patients With Crohn’s Disease. Gastroenterology 154, 2097–2110 (2018).

7. J. F. Tarlton et al., The role of up-regulated serine proteases and matrix metalloproteinases in the pathogenesis of a murine model of colitis. Am J Pathol 157, 1927–1935 (2000).

8. X. Wang et al., GPR34-mediated sensing of lysophosphatidylserine released by apoptotic neutrophils activates type 3 innate lymphoid cells to mediate tissue repair. Immunity 54, 1123–1136 e1128 (2021).

9. G. Zhou et al., CD177(+) neutrophils as functionally activated neutrophils negatively regulate IBD. Gut 67, 1052–1063 (2018).

10. E. L. Campbell et al., Transmigrating neutrophils shape the mucosal microenvironment through localized oxygen depletion to influence resolution of inflammation. Immunity 40, 66–77 (2014).

11. R. Sumagin et al., Neutrophil interactions with epithelial-expressed ICAM-1 enhances intestinal mucosal wound healing. Mucosal Immunol 9, 1151–1162 (2016).

12. M. Jeziorska, N. Haboubi, P. Schofield, D. E. Woolley, Distribution and activation of eosinophils in inflammatory bowel disease using an improved immunohistochemical technique. J Pathol 194, 484–492 (2001).

13. Y. Raab, K. Fredens, B. Gerdin, R. Hallgren, Eosinophil activation in ulcerative colitis: studies on mucosal release and localization of eosinophil granule constituents. Dig Dis Sci 43, 1061–1070 (1998).

14. O. Saitoh et al., Fecal eosinophil granule-derived proteins reflect disease activity in inflammatory bowel disease. Am J Gastroenterol 94, 3513–3520 (1999).

15. M. Carlson, Y. Raab, C. Peterson, R. Hallgren, P. Venge, Increased intraluminal release of eosinophil granule proteins EPO, ECP, EPX, and cytokines in ulcerative colitis and proctitis in segmental perfusion. Am J Gastroenterol 94, 1876–1883 (1999).

16. G. Katinios et al., Increased Colonic Epithelial Permeability and Mucosal Eosinophilia in Ulcerative Colitis in Remission Compared With Irritable Bowel Syndrome and Health. Inflamm Bowel Dis 26, 974–984 (2020).

17. M. Lampinen et al., Eosinophil granulocytes are activated during the remission phase of ulcerative colitis. Gut 54, 1714–1720 (2005).

18. E. C. L. Wong et al., Improvement in serum eosinophilia is observed in clinical responders to ustekinumab but not adalimumab in inflammatory bowel disease. J Crohns Colitis 19 (2025).

19. T. Alhmoud et al., Outcomes of inflammatory bowel disease in patients with eosinophil-predominant colonic inflammation. BMJ Open Gastroenterol 7, e000373 (2020).

20. Z. Wang et al., Role of eosinophils in a murine model of inflammatory bowel disease. Biochem Biophys Res Commun 511, 99–104 (2019).

21. I. Jacobs et al., Eosinophil Depletion as a Potential Therapeutic Strategy in Acute and Chronic Intestinal Inflammation Based on a Dextran Sulfate Sodium Colitis Model. Inflamm Bowel Dis 31, 169–177 (2025).

22. J. C. Masterson et al., Eosinophil-mediated signalling attenuates inflammatory responses in experimental colitis. Gut 64, 1236–1247 (2015).

23. G. F. Sonnenberg, L. A. Fouser, D. Artis, Border patrol: regulation of immunity, inflammation and tissue homeostasis at barrier surfaces by IL-22. Nat Immunol 12, 383–390 (2011).

24. Q. Zhu et al., Epithelial dysfunction is prevented by IL-22 treatment in a Citrobacter rodentium-induced colitis model that shares similarities with inflammatory bowel disease. Mucosal Immunol 15, 1338–1349 (2022).

25. L. A. Zenewicz et al., Innate and adaptive interleukin-22 protects mice from inflammatory bowel disease. Immunity 29, 947–957 (2008).

26. C. L. Zindl et al., A nonredundant role for T cell-derived interleukin 22 in antibacterial defense of colonic crypts. Immunity 55, 494–511 e411 (2022).

27. M. Moniruzzaman, R. Wang, V. Jeet, M. A. McGuckin, S. Z. Hasnain, Interleukin (IL)-22 from IL-20 Subfamily of Cytokines Induces Colonic Epithelial Cell Proliferation Predominantly through ERK1/2 Pathway. Int J Mol Sci 20 (2019).

28. G. Pickert et al., STAT3 links IL-22 signaling in intestinal epithelial cells to mucosal wound healing. J Exp Med 206, 1465–1472 (2009).

29. K. Sugimoto et al., IL-22 ameliorates intestinal inflammation in a mouse model of ulcerative colitis. J Clin Invest 118, 534–544 (2008).

30. S. J. Aujla et al., IL-22 mediates mucosal host defense against Gram-negative bacterial pneumonia. Nat Med 14, 275–281 (2008).

31. Y. Zheng et al., Interleukin-22, a T(H)17 cytokine, mediates IL-23-induced dermal inflammation and acanthosis. Nature 445, 648–651 (2007).

32. S. C. Liang et al., Interleukin (IL)-22 and IL-17 are coexpressed by Th17 cells and cooperatively enhance expression of antimicrobial peptides. J Exp Med 203, 2271–2279 (2006).

33. J. Qiu et al., The aryl hydrocarbon receptor regulates gut immunity through modulation of innate lymphoid cells. Immunity 36, 92–104 (2012).

34. N. K. Crellin et al., Regulation of cytokine secretion in human CD127(+) LTi-like innate lymphoid cells by Toll-like receptor 2. Immunity 33, 752–764 (2010).

35. T. Glatzer et al., RORgammat(+) innate lymphoid cells acquire a proinflammatory program upon engagement of the activating receptor NKp44. Immunity 38, 1223–1235 (2013).

36. J. J. Lee, E. A. Jacobsen, M. P. McGarry, R. P. Schleimer, N. A. Lee, Eosinophils in health and disease: the LIAR hypothesis. Clin Exp Allergy 40, 563–575 (2010).

37. Y. S. Guo et al., Gastrin stimulates cyclooxygenase-2 expression in intestinal epithelial cells through multiple signaling pathways. Evidence for involvement of ERK5 kinase and transactivation of the epidermal growth factor receptor. J Biol Chem 277, 48755–48763 (2002).

38. Singer, II et al., Cyclooxygenase 2 is induced in colonic epithelial cells in inflammatory bowel disease. Gastroenterology 115, 297–306 (1998).

39. D. Meriwether et al., Macrophage COX2 Mediates Efferocytosis, Resolution Reprogramming, and Intestinal Epithelial Repair. Cell Mol Gastroenterol Hepatol 13, 1095–1120 (2022).

40. T. Inaba et al., Induction of cyclooxygenase-2 in monocyte/macrophage by mucins secreted from colon cancer cells. Proc Natl Acad Sci U S A 100, 2736–2741 (2003).

41. T. O. Ishikawa, M. Oshima, H. R. Herschman, Cox-2 deletion in myeloid and endothelial cells, but not in epithelial cells, exacerbates murine colitis. Carcinogenesis 32, 417–426 (2011).

42. M. Sonoshita, K. Takaku, M. Oshima, K. Sugihara, M. M. Taketo, Cyclooxygenase-2 expression in fibroblasts and endothelial cells of intestinal polyps. Cancer Res 62, 6846–6849 (2002).

43. J. Travers, M. E. Rothenberg, Eosinophils in mucosal immune responses. Mucosal Immunol 8, 464–475 (2015).

44. E. A. Jacobsen, A. G. Taranova, N. A. Lee, J. J. Lee, Eosinophils: singularly destructive effector cells or purveyors of immunoregulation? J Allergy Clin Immunol 119, 1313–1320 (2007).

45. R. Sugawara et al., Small intestinal eosinophils regulate Th17 cells by producing IL-1 receptor antagonist. J Exp Med 213, 555–567 (2016).

46. C. Neufert et al., Activation of epithelial STAT3 regulates intestinal homeostasis. Cell Cycle 9, 652–655 (2010).

47. C. A. Rouzer, L. J. Marnett, Cyclooxygenases: structural and functional insights. J Lipid Res 50 Suppl, S29–34 (2009).

48. B. Martin, K. Hirota, D. J. Cua, B. Stockinger, M. Veldhoen, Interleukin-17-producing gammadelta T cells selectively expand in response to pathogen products and environmental signals. Immunity 31, 321–330 (2009).

49. S. Trifari, C. D. Kaplan, E. H. Tran, N. K. Crellin, H. Spits, Identification of a human helper T cell population that has abundant production of interleukin 22 and is distinct from T(H)-17, T(H)1 and T(H)2 cells. Nat Immunol 10, 864–871 (2009).

50. M. Cella et al., A human natural killer cell subset provides an innate source of IL-22 for mucosal immunity. Nature 457, 722–725 (2009).

51. Y. Chung et al., Expression and regulation of IL-22 in the IL-17-producing CD4+ T lymphocytes. Cell Res 16, 902–907 (2006).

52. W. Zhou et al., ZBTB46 defines and regulates ILC3s that protect the intestine. Nature 609, 159–165 (2022).

53. M. Fukata et al., Cox-2 is regulated by Toll-like receptor-4 (TLR4) signaling: Role in proliferation and apoptosis in the intestine. Gastroenterology 131, 862–877 (2006).

54. S. L. Brown et al., Myd88-dependent positioning of Ptgs2-expressing stromal cells maintains colonic epithelial proliferation during injury. J Clin Invest 117, 258–269 (2007).

55. A. Gurtner et al., Active eosinophils regulate host defence and immune responses in colitis. Nature 615, 151–157 (2023).

56. T. Grosser, S. Fries, G. A. FitzGerald, Biological basis for the cardiovascular consequences of COX-2 inhibition: therapeutic challenges and opportunities. J Clin Invest 116, 4–15 (2006).

57. K. Kabashima et al., The prostaglandin receptor EP4 suppresses colitis, mucosal damage and CD4 cell activation in the gut. J Clin Invest 109, 883–893 (2002).

58. D. G. Ribaldone et al., Coxib’s Safety in Patients with Inflammatory Bowel Diseases: A Meta-analysis. Pain Physician 18, 599–607 (2015).

59. Y. El Miedany, S. Youssef, I. Ahmed, M. El Gaafary, The gastrointestinal safety and effect on disease activity of etoricoxib, a selective cox-2 inhibitor in inflammatory bowel diseases. Am J Gastroenterol 101, 311–317 (2006).

60. R. Matuk et al., The spectrum of gastrointestinal toxicity and effect on disease activity of selective cyclooxygenase-2 inhibitors in patients with inflammatory bowel disease. Inflamm Bowel Dis 10, 352–356 (2004).

61. U. Mahadevan, E. V. Loftus, Jr., W. J. Tremaine, W. J. Sandborn, Safety of selective cyclooxygenase-2 inhibitors in inflammatory bowel disease. Am J Gastroenterol 97, 910–914 (2002).

62. G. F. Bonner, Exacerbation of inflammatory bowel disease associated with use of celecoxib. Am J Gastroenterol 96, 1306–1308 (2001).

63. M. J. Docherty, R. C. Jones, 3rd, M. S. Wallace, Managing pain in inflammatory bowel disease. Gastroenterol Hepatol (N Y) 7, 592–601 (2011).

64. L. Maiden et al., Long-term effects of nonsteroidal anti-inflammatory drugs and cyclooxygenase-2 selective agents on the small bowel: a cross-sectional capsule enteroscopy study. Clin Gastroenterol Hepatol 5, 1040–1045 (2007).

65. R. Duffin et al., Prostaglandin E(2) constrains systemic inflammation through an innate lymphoid cell-IL-22 axis. Science 351, 1333–1338 (2016).

66. L. Zhou et al., Group 3 innate lymphoid cells produce the growth factor HB-EGF to protect the intestine from TNF-mediated inflammation. Nat Immunol 23, 251–261 (2022).

67. Y. R. Na et al., Prostaglandin E(2) receptor PTGER4-expressing macrophages promote intestinal epithelial barrier regeneration upon inflammation. Gut 70, 2249–2260 (2021).

68. Y. Li et al., COX-2-PGE(2) Signaling Impairs Intestinal Epithelial Regeneration and Associates with TNF Inhibitor Responsiveness in Ulcerative Colitis. EBioMedicine 36, 497–507 (2018).

69. L. Xu et al., Eosinophils protect against acetaminophen-induced liver injury through cyclooxygenase-mediated IL-4/IL-13 production. Hepatology 77, 456–465 (2023).

70. L. A. Dieleman et al., Chronic experimental colitis induced by dextran sulphate sodium (DSS) is characterized by Th1 and Th2 cytokines. Clin Exp Immunol 114, 385–391 (1998).

71. C. Morral et al., Isolation of Epithelial and Stromal Cells from Colon Tissues in Homeostasis and Under Inflammatory Conditions. Bio Protoc 13, e4825 (2023).

72. C. Ibeakanma, S. Vanner, TNFalpha is a key mediator of the pronociceptive effects of mucosal supernatant from human ulcerative colitis on colonic DRG neurons. Gut 59, 612–621 (2010).

