## Supplementary Methods, Figures S1 to S6, Tables S1 and SI References for "Eosinophil-derived COX-2 protects against experimental colitis through the PGE_2_-IL-22 axis"

**This PDF file includes:**

Supplementary Methods
Figures S1 to S6
Tables S1
SI References

**Supporting Information**

***Supplementary Methods***

**Experimental colitis mouse model**

DSS or TNBS-induced colitis models were performed in 10- to 12-week-old male mice**.** For DSS-colitis, the mice were administered with 3% (w/v) DSS (Thermo Scientific, #9011-18-1, 40,000) in their drinking water for 5 consecutive days and then were switched to sterile water for another 4 days(1). Mice were monitored daily for weight and disease activity index (DAI). DAI considered stool consistency (0-4), presence of blood (0-4), and weight loss (0-4). The scores were added with a maximum possible score of 12. Mice were sacrificed on day 6 or day 9 to measure colon length and collect colon tissues for further analysis. For TNBS-colitis, epicutaneous skin sensitization with the hapten, tri-nitro-benzosulfonic acid (TNBS) (1% v/v; EtOH; Sigma Chemical) was performed under anesthesia on experimental day -7 in mice. On experimental day 0, anesthetized mice received the challenge of intra-rectal TNBS (5ul/g; 2.5% v/v; 40% EtOH) using a 20-gauge soft tygon plastic catheter. Vehicle control animals received a corresponding volume of 40% (v/v) ethanol alone(2). Body weight loss and DAI were assessed daily. Mice were sacrificed on day 3 to measure colon length and collect colon tissues for further analysis. For exogenous IL-22 treatment, mice were i.p. injected with rmIL-22 (R&D, #582-ML, 500 ng/mouse) on days 2, 5, 8 of DSS treatment. Control mice received a similar volume of Phosphate Buffered Saline (PBS). 16,16-dimethyl prostaglandin $E_2$ (Caymen, #14750, $dmPGE_2$) dissolved in methyl acetate. After methyl acetate was evaporated under a nitrogen stream, $dmPGE_2$ was immediately dissolved in nitrogen-purged ethanol and kept as a stock solution at a concentration of 0.5 mg/ml. The stock $dmPGE_2$ solution was diluted with saline immediately before use and kept on ice. For exogenous $PGE_2$ treatment, mice were i.p. injected with $dmPGE_2$ (10 μg/kg) on days 1, 3, 5, 8 of DSS treatment. Control mice received a similar volume of saline.

**Histology and Immunohistochemistry.**

In detail, four independent parameters were assessed, including inflammation (0-3), extent of injury (0-3), crypt damage (0-4), and regeneration (0-4). The score for each parameter was then multiplied by a factor reflecting the percentage of tissue involvement and combined for a maximum possible score of 40. IHC staining was performed using paraffin-embedded sections to determine the expression of MBP in the colon. Briefly, endogenous peroxidases were inactivated by 3% hydrogen peroxide. Nonspecific signals were blocked using 2.5% goat serum. Mouse-anti-EPX (Clone: MM25 82.2.2) and rat anti-MBP (Clone: MT2 14.7.3) from Dr. Elizabeth Jacobsen (Mayo Clinic Arizona) were used, and its staining was performed as previously described(3, 4). The slides were counterstained with hematoxylin and then washed and mounted with a mounting medium. Histologic images were captured on a Zeiss microscope.

**RNAscope analysis**

RNAscope™ Probe-Mm-Ptgs2 (#316621) was used in this study. TSA Vivid fluorophore 570 (#323272) was used for detection. Following RNAscope signal amplification, colon sections were blocked in 5% Donkey serum prior to incubation with Mouse-anti-EPX (Clone: MM25 82.2.2). A secondary antibody (Alexa Fluor 488 Donkey Anti-mouse, Biolegend) and DAPI were then applied to section before mounting. Imaging was captured using a Leica SP5 confocal microscope.

**Isolation of colonic lamina propria cells (LPCs)**

Colon tissue was excised, adipose tissue, mesentery, and mesenteric lymph nodes were carefully removed. Intestinal lumens cleaned and incubated in $Ca^{2+}/Mg^{2+}$-free HBSS (2% FCS) with 2.5 mM EDTA at 37°C to release IECs by gentle shaking. The supernatants were removed. The remaining colon fragments were collected and were digested with collagenase VIII (1.5 mg/mL) and DNase I (50 µg/mL) in $Ca^{2+}/Mg^{2+}$-free HBSS (2% FCS) at 37°C. The digested tissue was filtered, washed, and pelleted by centrifugation. Single-cell suspensions were stained with fluorochrome-conjugated antibodies and analyzed by flow cytometry following standard gating and viability exclusion.

**Flow cytometry analysis**

Cell surface staining was performed by incubating cells ($1x10^6$/per tube) with antibodies for 30 mins at 4°C after blocking with anti-CD16/CD32 (#101302, Biolegend). Dead cells were excluded by staining with blue fluorescent reactive dye (1:200, Invitrogen, 2176884). Flurochrome-conjugated antibodies against CD45 (1:200, clone: 30-F11), CD11b (1:200, clone: M1/70), Siglec-F (1:200, clone: E50-2440), CCR3 (1:200, clone: JO73E5), CD90.2(1:200, clone:30-H12), and CCR6 (1:200, Clone:29-2L17) were purchased from eBioscience or BD Biosciences or Biolegend. For IL-17A, RORrt and IL-22 intracellular staining, LPCs were treated with 50 ng/ml phorbol-12-myristate 13-acetate (PMA), 2.5 mg/ml monensin and 1 mg/ml ionomycin for 4 hours at 37C, 5% $CO_2$, before staining with anti-IL-17A (1:100, Clone:TC1118H10.1), anti-RORγt (1:100, clone: Q31-378) and anti-IL-

22 (1:100, clone: JOP22) antibody using a Fixation & Permeabilization Kit (Invitrogen). The cells were analyzed using a CytoFLEX LX flow cytometer (Beckman Coulter) and the data were analyzed with FlowJo v10.0.7.

**RNA isolation and quantitative real-time polymerase chain reaction (qRT-PCR)**

Total RNA was purified from colon tissues or culture cells using Pure-Link RNA Kit (GeneDopt) according to the manufacturer's protocol, and 1 μg RNA was reverse transcribed into cDNA using an iScript cDNA Synthesis Kit (Bio-Rad). Relative quantitative gene expression was measured with SYBR Green PCR Supermix from GenDEPOT. 18S rRNA was used as an internal standard. The primers used for qRT-PCR are listed in *Table S1*.

***Ex vivo* colon cultures and Enzyme-linked immunosorbent assay (ELISA)**

Colon tissues were cultured ex vivo as previously described with minor modifications(5). Briefly, colons were flushed with ice-cold PBS, opened longitudinally, and cut into ~100 mg segments. Tissue sections were incubated in 0.5 mL RPMI 1640 medium supplemented with 10% FBS, 100 IU/mL penicillin, 100 μg/mL streptomycin, 1 mM sodium pyruvate, and 2 mM L-glutamine in 24-well plates at 37°C with 5% $CO_2$ for 48 hours. Culture supernatants were collected, and IL-22 levels were measured by ELISA (Biolegend, 436304) following the manufacturer’s instructions.

### Figures

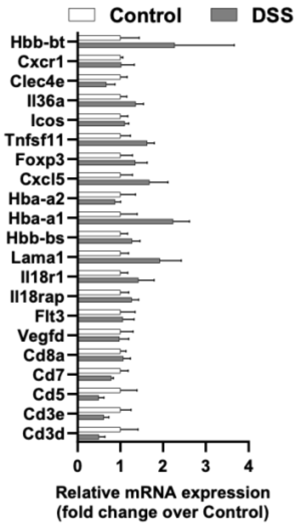


***Fig.S1. Validation of DGEs mRNA levels.*** WT C57Bl/6 mice were treated with regular water or 3% DSS for 5 days. On day 6, colonic eosinophils were isolated/purified by MACS and subjected to qPCR analyses. (n=3/group). mRNA levels of 21 genes in colonic eosinophils were measured. A two-tailed unpaired Student's t test with Welch's correction was performed.

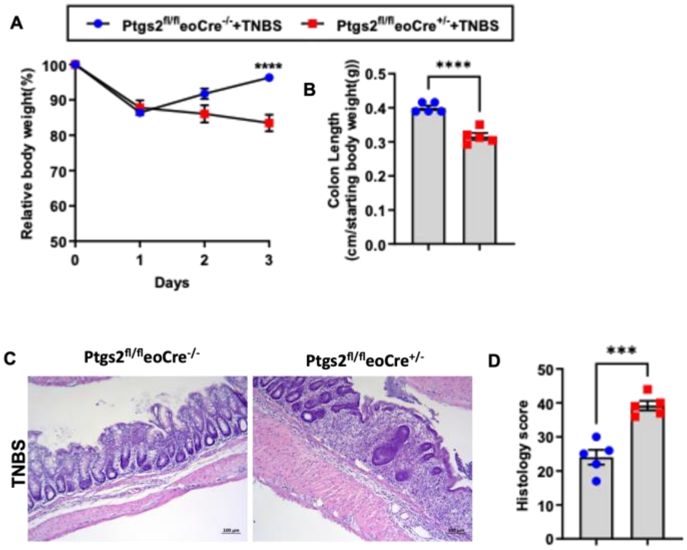


***Fig.S2. Deletion of Ptgs2 in eosinophils develop more severe TNBS-induced colitis***. (A-D) Ptgs2$^{fl/fl}$eoCre$^{+/-}$ mice and their WT littermates were treated with 2.5% TNBS, and colon tissues were collected on day 3. (n=5/group) (A) Body weight changes and (B) colon lengths were measured. (C-D) Representative colon histology and histological scores. Data are means ± SEM. In all scatter plots with bars, each data point represents one individual. A two-tailed unpaired Student's t test with Welch's correction was performed in B and D. One-way ANOVA was performed in A. ***p < 0.001, ****p < 0.0001.

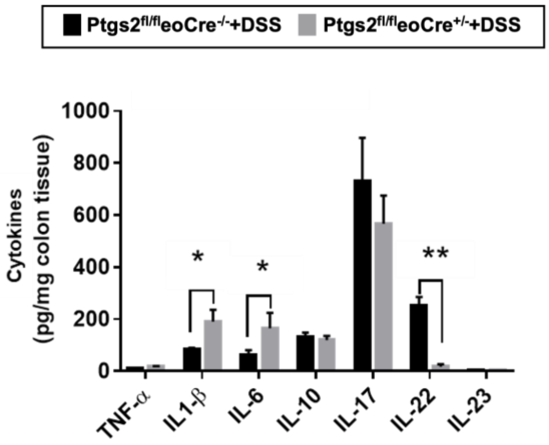


***Fig.S3. Compare colon tissue cytokines between Ptgs2$^{fl/fl}$eoCre$^{+/-}$ and their WT littermates during colitis.*** Ptgs2$^{fl/fl}$eoCre$^{+/-}$ mice and their WT littermates were treated with 3% DSS for 5 days. Colon tissues were collected on day 6. Protein levels of colitis-associated cytokines were measured in the supernatant (n=3/group). One-way ANOVA was performed in A. *$p < 0.05$, **$p < 0.001$.

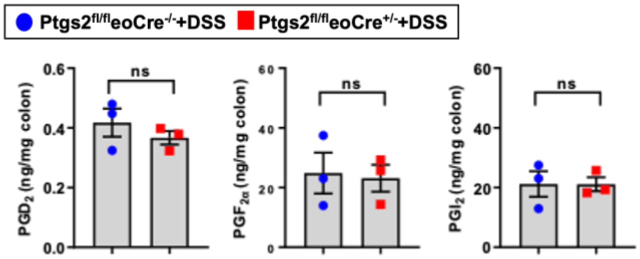


***Fig.S4. Colonic levels of PGs after DSS treatment.*** Ptgs2$^{fl/fl}$eoCre$^{+/-}$ mice and their WT littermates were treated with 3% DSS for 5 days and collected colon tissues on day 6. Colonic $PGD_2$, $PGF_{2\alpha}$, and $PGI_2$ levels were measured by ELISA (n=3/group). Data are means ± SEM. In all scatter plots with bars, each data point represents one individual. A two-tailed unpaired Student's t test with Welch's correction was performed.

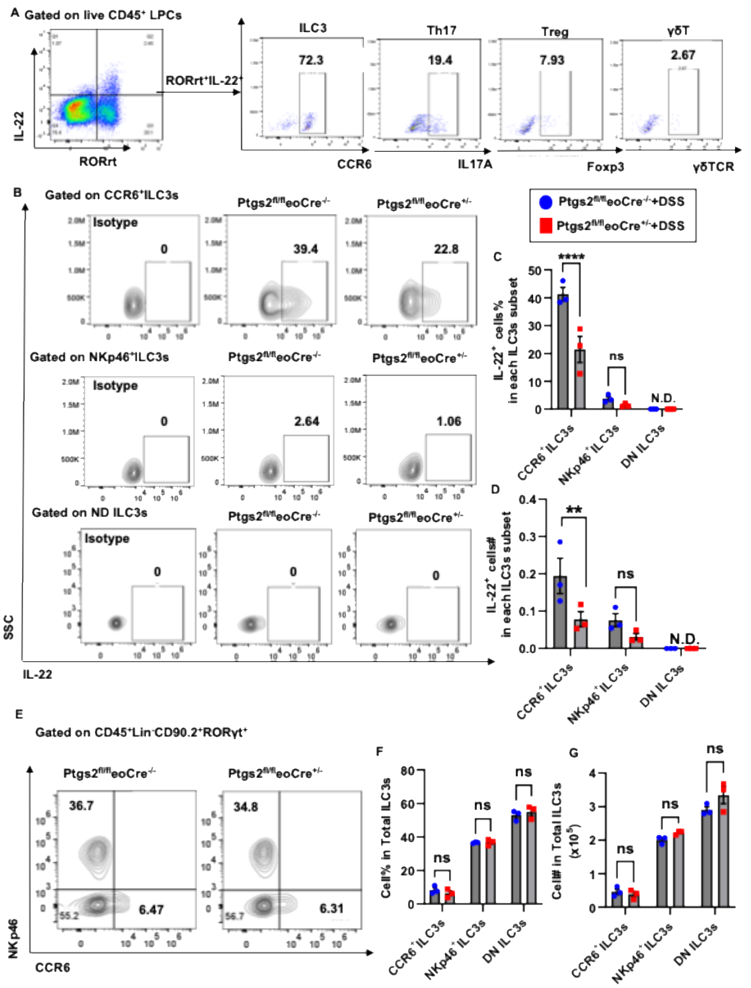


***Fig.S5 CCR6$^+$ILC3s are the major cellular source of IL-22 in the colon of DSS-treated WT mice.*** (A)WT C57Bl/6 mice were treated with DSS for 5 days. On day 6, LPs were isolated and stained for intracellular IL-22. The IL-22-positive cells are gated, and the proportion of ILC3s (CCR6$^+$) that express IL-22 among total CD45$^+$CD90.2$^+$RORrt$^+$Lin$^-$IL-22$^+$ is shown. (B-E) Ptgs2$^{fl/fl}$eoCre$^{+/-}$ mice and their WT littermates were treated with 3% DSS for 5 days. On day 6, colon lamina propria cells (LPCs) were isolated and divided into two groups. (B-D) Representative flow cytometry, the percentage and numbers of IL-22$^+$ILC3s subsets in the LPs were measured and quantified. (D-F) Representative flow cytometry, the percentage and numbers of ILC3 subsets in the LPCs were measured and quantified. Data are means ± SEM. In all scatter plots with bars, each data point represents one individual. Two-way ANOVA was performed in C-D and F-G. *$p < 0.05$, **$p < 0.01$, ***$p < 0.001$. All experiments were repeated a minimum of three times.

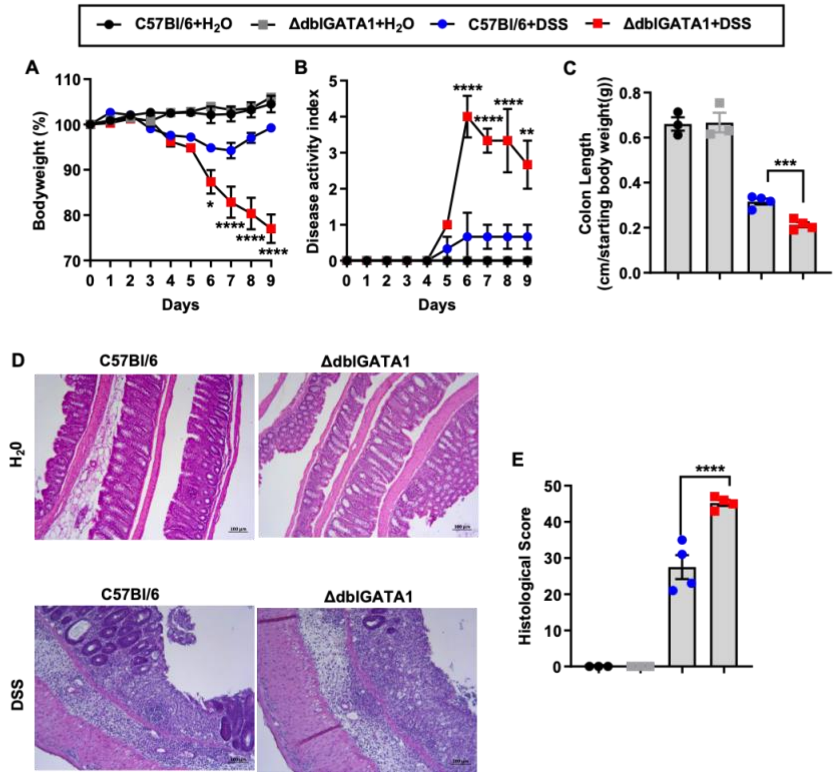


***Fig.S6. Eosinophils deficiency exhibits more severe colitis***. ΔdblGATA1 mice and their WT counterparts were treated with 3% DSS for 5 day and allowed to recover for 4 days before collection of colon tissues on day 9. (n=4/group) Control mice were treated with regular water. (n=3/group) (A) Body weight changes (B) disease activity index and (C) colon lengths were measured. (D-E) Representative colon histology and histological scores. Data are means ± SEM. In all scatter plots with bars, each data point represents one individual. Two-way ANOVA was performed A, B, C and E. *$p < 0.05$, **$p < 0.01$, ***$p < 0.001$, ****$p < 0.0001$

**Table S1. The primers used for qRT-PCR**

| Genes (Mouse) | | *5'-3'* |
|---|---|---|
| Ptgs2 | *Forward* | *TGAGCAACTATTCCAAACCAGC* |
| | *Reverse* | *GCACGTAGTCTTCGATCACTATC* |
| 18s | *Forward* | *ACGGAAGGGCACCACCAGGA* |
| | *Reverse* | *CACCACCACCCACGGAATCG* |
| Csf3 | *Forward* | *ATGGCTCAACTTTCTGCCCAG* |
| | *Reverse* | *CTGACAGTGACCAGGGGAAC* |
| Il11 | *Forward* | *TGTTCTCCTAACCCGATCCCT* |
| | *Reverse* | *CAGGAAGCTGCAAAGATCCCA* |
| Lcn2 | *Forward* | *TGGCCCTGAGTGTCATGTG* |
| | *Reverse* | *CTCTTGTAGCTCATAGATGGTGC* |
| Mmp3 | *Forward* | *ACATGGAGACTTTGTCCCTTTTG* |
| | *Reverse* | *TTGGCTGAGTGGTAGAGTCCC* |
| Cxcl2 | *Forward* | *CCAACCACCAGGCTACAGG* |
| | *Reverse* | *GCGTCACACTCAAGCTCTG* |
| Tnfrsf9 | *Forward* | *CGTGCAGAACTCCTGTGATAAC* |

| | | |
|---|---|---|
| | *Reverse* | *GTCCACCTATGCTGGAGAAGG* |
| Clec4d | *Forward* | *ACCCGACATCCCCAACTGAT* |
| | *Reverse* | *CTCTCGTCCAGCGTAAAAAGT* |
| S100a9 | *Forward* | *ATACTCTAGGAAGGAAGGACACC* |
| | *Reverse* | *TCCATGATGTCATTTATGAGGGC* |
| S100a8 | *Forward* | *AAATCACCATGCCCTCTACAAG* |
| | *Reverse* | *CCCACTTTTATCACCATCGCAA* |
| Nlrp3 | *Forward* | *ATTACCCGCCCGAGAAAGG* |
| | *Reverse* | *TCGCAGCAAAGATCCACACAG* |
| Arg2 | *Forward* | *TCCTCCACGGGCAAATTCC* |
| | *Reverse* | *GCTGGACCATATTCCACTCCTA* |
| Arg1 | *Forward* | *CTCCAAGCCAAAGTCCTTAGAG* |
| | *Reverse* | *AGGAGCTGTCATTAGGGACATC* |
| Il1r2 | *Forward* | *GTTTCTGCTTTCACCACTCCA* |
| | *Reverse* | *GAGTCCAATTTACTCCAGGTCAG* |
| Csf3r | *Forward* | *CTGATCTTCTTGCTACTCCCCA* |
| | *Reverse* | *GGTGTAGTTCAAGTGAGGCAG* |
| Ctsg | *Forward* | *AGGGTTTCTGGTGCGAGAAG* |
| | *Reverse* | *GTTCTGCGGATTGTAATCAGGAT* |
| Il6 | *Forward* | *TAGTCCTTCCTACCCCAATTTCC* |
| | *Reverse* | *TTGGTCCTTAGCCACTCCTTC* |
| Il23a | *Forward* | *ATGCTGGATTGCAGAGCAGTA* |
| | *Reverse* | *ACGGGGCACATTATTTTTAGTCT* |
| Il22 | *Forward* | *ATGAGTTTTTCCCTTATGGGGAC* |
| | *Reverse* | *GCTGGAAGTTGGACACCTCAA* |

### SI References


1. P. Smith *et al.*, Infection with a helminth parasite prevents experimental colitis via a macrophage-mediated mechanism. *J Immunol* **178**, 4557-4566 (2007).
2. A. K. Czopik *et al.*, HIF-2alpha-dependent induction of miR-29a restrains T(H)1 activity during T cell dependent colitis. *Nat Commun* **15**, 8042 (2024).
3. S. I. Ochkur *et al.*, Cys-leukotrienes promote fibrosis in a mouse model of eosinophil-mediated respiratory inflammation. *Am J Respir Cell Mol Biol* **49**, 1074-1084 (2013).
4. S. I. Ochkur *et al.*, Coexpression of IL-5 and eotaxin-2 in mice creates an eosinophil-dependent model of respiratory inflammation with characteristics of severe asthma. *J Immunol* **178**, 7879-7889 (2007).
5. C. Ibeakanma, S. Vanner, TNFalpha is a key mediator of the pronociceptive effects of mucosal supernatant from human ulcerative colitis on colonic DRG neurons. *Gut* **59**, 612-621 (2010).
